# The roles and synthesis of inorganic polyphosphate in *Bacillus cereus*

**DOI:** 10.64898/2026.08.04.742738

**Authors:** Christine Kim, Lilla Fournier, Michael J. Gray, Christopher W. Hamm

## Abstract

Inorganic polyphosphate (polyP) is a universally conserved biopolymer central to bacterial stress survival, yet understanding of its roles derives almost entirely from Gram-negative models in which polyP accumulates intracellularly following nutrient downshift. We examined polyP metabolism in the Gram-positive spore-forming bacterium *Bacillus cereus* using deletions of the polyP kinases PPK1 and PPK2 and the exopolyphosphatase PPX. Intracellular polyP synthesis required PPK1 and was opposed by PPX and PPK2: *ppx* mutants accumulated polyP in sporulation medium by 24 hours, *ppx ppk2* double mutants accumulated more, and no *ppk1* mutant accumulated any. A *ppk1 ppx* double mutant could not be generated, suggesting that unopposed PPK2 activity is lethal. Unlike *Escherichia coli* and *Pseudomonas aeruginosa*, *B. cereus* did not accumulate polyP after shift to minimal medium, increasing only modestly in stationary phase. Fluorescence and transmission electron microscopy localized intracellular polyP to electron-dense granules within ribosome-depleted cytoplasm. Cells bearing these granules remained membrane-intact yet failed to resume growth over 8 hours in rich medium, leading us to propose that polyP drives ribosome sequestration into condensates and a hibernation-like state. Unexpectedly, *B. cereus* also released close to 100µM polyP extracellularly during late stationary phase, even in a *ppk1 ppk2* mutant lacking both known synthetases. Extracellular polyP resisted hydrolysis by purified PPX even after deproteinization, indicating an atypical structure. *Bacillus thuringiensis* and *Bacillus anthracis* released similar amounts of extracellular polyP. Together these results identify two distinct polyP pools in the *B. cereus* group: a PPK1-dependent intracellular pool and an extracellular pool made by an uncharacterized pathway.

**Importance:** *Bacillus cereus* is a spore-forming bacterium that causes foodborne illness and persists in soil and food-processing environments, where survival depends on managing phosphate and energy reserves during starvation. Inorganic polyphosphate (polyP), an ancient polymer used by nearly all cells to withstand stress, has been studied almost entirely as a molecule stored inside bacteria. We show that *Bacillus cereus* maintains two separate polyP pools. The internal pool is made by a known enzyme (PPK1) and is associated with dormant cells whose protein-making machinery appears to be packed away. The external pool is made without any known polyP-synthesizing enzyme, pointing to a novel polyP synthesis pathway that is shared with the close relatives *Bacillus thuringiensis* and *Bacillus anthracis*.

## Introduction

*Bacillus cereus* is a Gram-positive, spore forming bacteria that is ubiquitous in the environment ^1–6^. Its ability to form environmentally resilient spores enables persistence throughout the food chain, where it is a common cause of foodborne illness ^1,3,7^. Because of its widespread distribution in soil and food products, *B. cereus* is frequently ingested and may transiently colonize the human gastrointestinal tract, where it can act as an opportunistic pathogen ^8,9^. Although most commonly associated with foodborne illness, *B. cereus* can also cause localized wound and ocular infections, as well as severe systemic disease that may be fatal ^10^. Members of the *B. cereus* group are also of considerable interest due to their close relationship with *Bacillus anthracis*, the causative agent of anthrax ^11^, and *Bacillus thuringiensis,* which is widely used as a biological pesticide worldwide ^10,12,13^. The ability of these organisms to persist in diverse environments and establish successful host interactions highlights the importance of understanding the molecular mechanisms that contribute to bacterial stress adaptation, survival, and pathogenesis in these species.

One such mechanism involves inorganic polyphosphate (polyP), a widely distributed biopolymer that is highly conserved across all domains of life ^14–18^. PolyP is a linear polymer of orthophosphates that can be made up of 10-1000 orthophosphates linked by high energy bonds ^19^. In bacteria, polyP participates in numerous cellular processes, including the modulation of gene expression during stress responses ^17,20–24^, metal chelation ^25–27^, energy and phosphate storage ^28–30^, protein stabilization through chaperone-like activity ^31,32^, and other functions. Although polyP is not essential for bacterial viability, its absence renders cells highly sensitive to environmental stressors, including antibiotics, oxidative damage, osmotic stress, and nutrient limitation ^14,15,26,29,33–46^. Notably, pathogenic bacteria deficient in polyP show severe attenuation of virulence and increased sensitivity to antibiotics ^47–54^, highlighting polyP as a central regulator of bacterial fitness and pathogenicity. These findings have identified polyP metabolism and regulation as an attractive target for antimicrobial drug development, particularly since mammals lack homologs of any known microbial polyP synthesis enzymes ^14,30,36,46,55–59^.

In prokaryotes, polyP is synthesized by two known types of polyphosphate kinases (PPK’s) with limited sequence homology to one another ^29,30,43,57,60,61^: PPK1 and PPK2, both of which are widespread among both Gram-negative and Gram-positive bacteria (**Fig 1**)^17,24,30,62,63^. PolyP is primarily degraded by exopolyphosphatase (PPX)^23,64,65^ (**Fig 1**). PPK1, first identified in *Escherichia coli* in the 1990’s, preferentially produces polyP from ATP, although it can also catalyze the reverse reaction (**Fig 1**)^63,66^. PPK2 enzymes, originally identified in *Pseudomonas aeruginosa*, preferentially catalyze polyP degradation to generate nucleoside phosphates, exhibiting approximately a 75-fold greater activity for degradation than for synthesis, despite retaining the capacity to synthesize polyP ^30,43,44,53,61,62,67^. PPK2s are further classified into three subfamilies based on their preference for a specific nucleoside phosphate substrate. Class I preferentially phosphorylates nucleoside diphosphates, Class II nucleoside monophosphates, and Class III can phosphorylate both nucleoside di- and monophosphates ^30,68^. While *Pseudomonas* contains PPK1 as well as all three types of PPK2 ^30^, the genome of *B. cereus* encodes only single PPK1, PPX, and class-II PPK2 homologs (**Fig. 1**)^30,54,64,68,69^.

**Figure 1.**
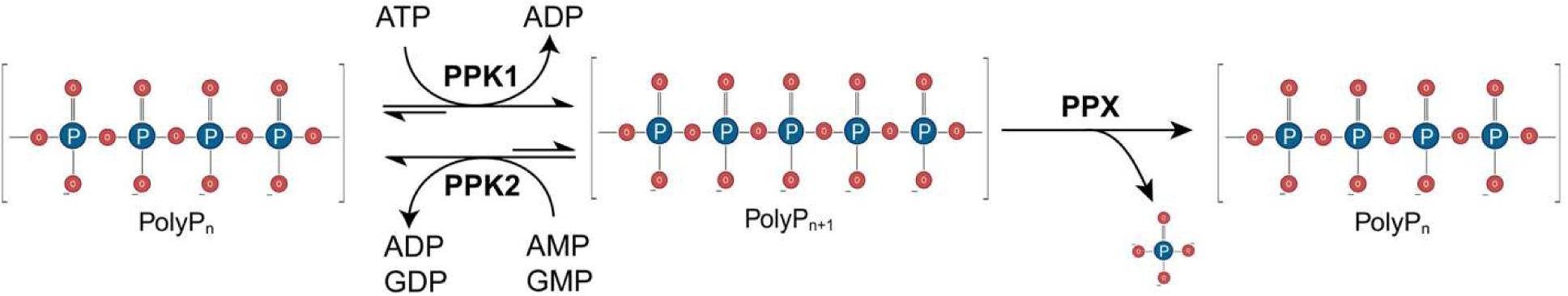
Schematic of known enzymes contributing to inorganic polyP synthesis and degradation in *Bacillus cereus*.

Despite being found in nearly all living organisms, almost all of the work on polyP in prokaryotes has been done in Gram-negative bacteria, specifically *E. coli* ^14,32,33,35,38,39,55^ and *P. aeruginosa* ^30,41,57,62,67,70^. However, in 2004, Arthur Kornberg’s lab established *B. cereus* as a model organism for studying polyP in Gram-positives ^69^. They reported that in *B. cereus* ATCC 14579, deletions of PPK1, PPK2, and PPX affected motility, biofilm formation and sporulation. There have been almost no studies into the molecular biology of Gram-positive bacterial polyP since. This means that, despite the established importance of polyP in bacterial physiology and pathogenesis, the mechanisms governing its synthesis, regulation, and biological functions in Gram-positive organisms remain poorly defined. This knowledge gap is particularly relevant in *B. cereus*, where understanding how polyP contributes to cellular fitness and host interaction may reveal novel insights into bacterial pathogenesis and identify new opportunities for antimicrobial development.

## Materials and Methods

### Strains, plasmids and growth conditions

The Strains and plasmids used in this study are listed below in Table 1.

**Table 1.** List of strains and plasmids used in this study.

| Name | Genotype | Source |
| --- | --- | --- |
| CH0082 | *B. cereus* ATCC 14579 | BGSC |
| CH0193 | *B. cereus* ATCC 14579 $\Delta ppk1$ | This study |
| CH0197 | *B. cereus* ATCC 14579 $\Delta spo0A$ | This study |
| CH0201 | *B. cereus* ATCC 14579 $\Delta ppx$ | This study |
| CH0212 | *B. cereus* ATCC 14579 $\Delta ppk2$ | This study |
| CH0217 | *B. cereus* ATCC 14579 $\Delta ppx \Delta ppk2$ | This study |
| CH0221 | *B. cereus* ATCC 14579 $\Delta ppk1 \Delta ppk2$ | This study |
| CH0244 | *B. cereus* ATCC 14579 $\Delta ppx \Delta ppk1 \Delta ppk2$ | This study |
| 4D1 | *Bacillus thuringiensis* (Kurstaki HD1) | Charles<br>Turnbough, <sup>71</sup> |
| Sterne<br>34F2 | *Bacillus anthracis* (USAMR11D) (avirulent isolate<br>lacking virulence plasmid pXO2) | Charles<br>Turnbough, <sup>72</sup> |
| *dam<sup>-</sup>/dcm<sup>-</sup>*<br>*E. coli* | *dam<sup>-</sup>/dcm<sup>-</sup> E. coli* chemically competent cells from New<br>England BioLabs, Ipswich, MA, USA, Cat#C2925H | NEB |
| **Plasmids** | **Genotype/purpose** | **Source** |
| pMiniMAD | Cloning vector for *Bacillus* species, *oriTsBs*-MLS<sup>R</sup> *Ec*-<br>Amp<sup>R</sup> | Gift from Dan<br>Kearns, <sup>73</sup> |
| pMiniMAD-<br>$\Delta ppk1$ | Deletion plasmid for *ppk1* in *B. cereus* constructed by<br>GenScript. | GenScript |
| pMiniMAD-<br>$\Delta ppx$ | Deletion plasmid for *ppx* in *B. cereus* constructed by<br>GenScript. | GenScript |
| pMiniMAD-<br>$\Delta ppk2$ | Deletion plasmid for *ppk2* in *B. cereus* constructed by<br>GenScript. | GenScript |
| pMiniMAD-<br>$\Delta spo0A$ | Deletion plasmid for *spo0A* in *B. cereus* constructed by<br>GenScript. | GenScript |
| pMiniMAD-<br>$\Delta ppx$<br>$\Delta ppk1$ | Deletion plasmid for the *ppx* and *ppk1* gene cluster in *B.*<br>*cereus* constructed by GenScript. | GenScript |

All strains were routinely grown in Lysogeny Broth (LB) medium (10 g/L tryptone, 5 g/L yeast extract, 5 g/L NaCl) or in Bacto^TM^ Tryptic Soy Broth (TSB) (Soybean-Casein Digest Medium) at 37°C in broth or on 1.5% Bacto agar as appropriate. When appropriate antibiotics were added (MLS: 0.5 μg/ml erythromycin and 2.5 μg/ml lincomycin, ampicillin 100 µg/ml) to select for markers. For sporulation media, 1/10^th^ TSB media was used (example: 10 ml of TSB with 90 ml of sterile H_2_O) and cells were grown at 30°C. Strains were always grown in sterile, 250 ml beveled flasks with shaking at 180 rpm to ensure proper oxygenation of cells. Markerless replacement for gene deletions was performed with the pMiniMAD vector (gift from Dan Kearns, Indiana University) for allelic replacement, as previously described ^73^. All mutant strains were confirmed with full genome sequencing through SeqCenter (Pittsburgh, PA) using Illumina Whole Genome Sequencing. The DNA sequences for the plasmids constructed in this study were codon-optimized and custom-synthesized, then cloned into the pMiniMAD cloning vector by GenScript (Piscataway, NJ, USA). All plasmid sequences were confirmed via whole plasmid sequencing using Oxford Nanopore Technology (Plasmidsaurus, LLC).

### Strain Construction

The markerless gene replacement in the strains below were added in succession using the pMiniMAD vector. A pMiniMAD vector containing our gene deletion of interest was synthesized by GenScript (Piscataway, NJ, USA) with 600 bp of genomic DNA sequence on either side of the gene of interest and propagated in *E. coli dcm^-^/dam^-^* for storage and propagation. Plasmids must be kept in a nonmethylated strain of *E. coli* as *B. cereus* will not uptake methylated DNA ^74^. The plasmid containing the desired gene deletion was directly transformed into *B. cereus* ATCC 14579 via electroporation ^75^ and selected on MLS (0.5 μg/ml erythromycin and 2.5 μg/ml lincomycin). Five to 10 transductants were then inoculated into liquid LB and kept in exponential phase at approximately 25°C for several hours to permit plasmid excision before being repeatedly diluted and grown in liquid LB at 37°C (restrictive for plasmid replication) to promote loss of excised plasmid. The cells were then plated, and single colonies were screened for the successful replacement by patching on plain LB and LB/MLS plates to verify plasmid loss, restreaked, verified by full genome sequencing at SeqCenter (Pittsburgh, PA) and stored at -80°C.

### PolyP Assay

Bacterial cultures were centrifuged at 16,100 x g for 2 minutes at room temperature to separate the cells from culture supernatants. JC-D7 (polyP binding dye) ^76,77^ was purchased from InvivoChem (Houston, TX, USA)(Cat. No. V22869) and dissolved in DMSO at 60 µM before dilution to 1 µM with 25 mM HEPES-KOH (pH 8.0). Extracellular polyP was quantified by combining 50 µl supernatants with 50 µl 1 µM JC-D7 in a black 96-well plate. Sodium polyP standards (0-100 µM, dissolved in 25 mM HEPES-KOH pH 8.0) (sodium polyP acquired from Acros Organics) were included for calibration on each plate. Following a 5-minute incubation at room temperature, fluorescence was measured in a Tecan Spark plate reader (Ex 405 nm/Em 535 nm).

Cell pellets were resuspended in 250 µL guanidinium thiocyanate (GITC) lysis buffer (4 M GITC, 50 mM Tris-HCl, pH 7)^39,78,79^ and disrupted by bead beating (2 x 40 s, 1 min interval, power level 6) in a Fisherbrand bead mill 24. Lysates were clarified by centrifugation at 16,100 x g for 2 min, and protein concentrations were determined using a Bradford assay with bovine serum albumin (BSA) standards. Intracellular polyP was isolated from lysates using silica membrane spin columns as previously described ^64^. Bound polyP was washed with 750 µL of 5 mM Tris-HCl pH 7.5, 50 mM NaCl, 5 mM EDTA, 50% EtOH, then eluted in 200 µL of 25 mM HEPES-KOH (pH 8.0) and quantified by mixing eluates (50 µl) with JC-D7 ^38,76,77,80^ dye (50 µl) in a black 96-well plate. After incubation for 5 min at room temperature, fluorescence was measured in a Tecan Spark plate reader (Ex 405 nm/Em 535 nm). PolyP concentrations were calculated from a polyP standard curve and normalized to total protein content, with results reported as nmol polyP mg^-1^ protein.

### Microscopy Imaging

Fluorescent and time-lapse microscopy was performed with a Keyence BZ-X810 all-in-one fluorescence microscope (Keyence Corporation, Osaka, Japan) equipped with a BZ Plan Apo Chromatic 60x oil objective. Images were captured using the BZ-X800 Viewer/Analyzer software and processed with BZ-X800 analyzer software or FIJI. Our microscope has a BZX DAPI filter, BZX TritC filter and BZX GFP Filter. We had a custom JC-D7 filter cube made by Chroma Technology, Bellows Falls, VT, USA which contains for excitation: Et 405/20x LP filter and T425LPL XR dichromatic mirror, and for emission: ET 450 LP filter. Automated time lapse microscopy and stage movement was controlled and run by BZ-X800 Software. Samples were maintained at 37°C using a Tokai Hit stage-top incubation system (Tokai Hit, Fujinomiya, Japan).

### Transmission Electron Microscopy Sample Preparation

Specimens were prepared for transmission electron microscopy (TEM) using a standard embedding protocol, where cell pellets were fixed in 2% glutaraldehyde prepared in 0.15 M sodium cacodylate (NaCaCo) buffer (pH 7.4) for a minimum of 1 hour. Following fixation, the samples were rinsed several times with sodium cacodylate buffer and post-fixed in 1% osmium tetroxide containing 0.8% potassium ferrocyanide in buffer for 1 hour. After additional buffer rinses for 1 hour, the samples were dehydrated through a graded ethanol series consisting of 50% ethanol for 10 minutes, 70% ethanol for 10 minutes, 95% ethanol for 10 minutes, and three changes of 100% ethanol for 15 minutes each. Samples were then infiltrated with two 15-minute changes of 100% propylene oxide followed by a 50:50 mixture of propylene oxide and Embed 812 epoxy resin (Electron Microscopy Sciences, Fort Washington, PA) for 12–18 hours. The samples were transferred through two fresh changes of 100% Embed 812 resin for at least 1 hour each before final embedding in fresh resin. Polymerization was performed at 60°C for 12–18 hours.

### Transmission Electron Microscopy Sectioning and Imaging

Polymerized resin blocks were initially sectioned at 0.5 µm using a diamond histology knife on an ultramicrotome. Thick sections were stained with Toluidine Blue and used to identify regions of interest for trimming prior to ultrathin sectioning. Ultrathin sections (70– 100 nm; silver to pale gold interference color) were cut with a diamond knife (Diatome; Electron Microscopy Sciences, Fort Washington, PA) and collected on copper mesh grids. Following drying, sections were contrasted with uranyl acetate and lead citrate. Grids were examined using a JEOL 1400 FLASH 120 kV transmission electron microscope (JEOL USA Inc., Peabody, MA). Digital images were acquired using an AMT NanoSprint43 Mark II camera (AMT Imaging, Woburn, MA) and transferred electronically via UAB BOX or other secure file-sharing methods.

### Neutral phenol/chloroform and ethanol precipitation of polyP

Culture supernatants were clarified by centrifugation (4,000 × g, 10 min, 4 °C) and passed through a 0.22 µm filter to remove residual cells. Five milliliters of cell-free supernatant were extracted with an equal volume of phenol:chloroform:isoamyl alcohol (25:24:1, pH 7.8–8.2); acid phenol was avoided owing to the instability of polyP under acidic conditions. Following centrifugation (4,000 × g, 10 min, room temperature), the aqueous phase was re-extracted with an equal volume of chloroform to remove residual phenol^81^. PolyP was precipitated from the recovered aqueous phase with 0.1 volume of 3 M ammonium acetate and 2.5 volumes of ice-cold ethanol at −20 °C overnight, pelleted at 10,000 × g for 30 min at 4 °C, washed once with 70% ethanol, air-dried briefly, and resuspended in 40 µL of 10 mM Tris-HCl (pH 7), 1 mM EDTA. Nuclease treatment was omitted, as DAPI staining discriminates polyP from co-purifying nucleic acids in-gel^81,82^.

Samples were mixed 1:1 with loading buffer (10 mM Tris-HCl pH 7, 1 mM EDTA, 30% glycerol, bromophenol blue) and resolved on 15.0% TBE-urea polyacrylamide gels (Invitrogen Cat#EC68855BOX) at 100 V for 1 h 45 min in 1× TBE. Gels were stained for 30 min in 20% methanol, 1% glycerol, 20 mM Tris base containing 20 µg/mL DAPI, destained for 60 min in the same solution lacking DAPI, and imaged. Gels were then exposed to 300 nm UV for 5 min and reimaged; under these conditions DAPI–polyP complexes preferentially photobleach, allowing polyP to be detected as negatively stained bands against a fluorescent nucleic acid background^82^. EDTA was maintained throughout to prevent DAPI staining of glycosaminoglycans.

### Exopolyphosphatase Treatment

ScPPX was purified as previously described ^32^. We used sodium polyP (Cas# 50813-16-6; average ∼ 45-mer) for standards. To 150 µl polyP sample solution and standard curve solutions of 0, 25, 50 and 100 µM potassium phosphate, we added 40 µl 5X ScPPX Reaction Buffer (100 mM Tris-HCl [pH 7.5], 25 mM MgCl_2_, 250 mM ammonium acetate), 9 µl dH_2_O, and 1 µl ScPPX (1 µg). We incubated the reactions for 15 minutes at 37°C, then added 50 µl of digested polyP sample to 50 µl of JC-D7, and determined polyP concentrations as described above.

### Data Availability

Genome sequences of all *B. cereus* strains used in this study have been deposited in the NIH Sequence Read Archive (accession number PRJNA1505627) and all other raw data are available on FigShare (10.6084/m9.figshare.c.8628284).

## Results

### Intracellular polyP accumulation

We first wanted to characterize the production of polyP in *B. cereus* using modern genetic tools and detection assays unavailable in the previous study of *B. cereus* polyP from 2004 ^69,76,77^. We used allelic exchange mutagenesis to individually delete *ppk1*, *ppk2*, *ppx*, and the global cell fate regulator *spo0A*^83,84^ as well as constructing double Δ*ppk1 Δppk2*, Δ*ppk2 Δppx*, and triple Δ*ppk1 Δppk2 Δppx* mutants. Surprisingly, despite multiple attempts, we were unable to make a Δ*ppx Δppk1* mutant, indicating that the presence of PPK2 alone appears to be lethal to *B. cereus*. To test the amount of polyP made by cells that are entering starvation, we grew cells in rich TSB media at 37°C and assayed the amount of polyP associated with the cells over a period of 72 hours (**Fig 2A**). We found little to no polyP accumulating in any strain of *B. cereus* over the course of 72 hours in TSB medium (**Fig 2A**). When we tested cells in sporulation media (1:10 TSB) grown at 30°C over the same time period, however, we found that cells lacking *ppx* were able to start accumulating detectable polyP as early as 24 hours (**Fig 2B**). The *ppx* mutant continued to accumulate polyP over 48 hours, but returned to wild-type levels at 72 hours (**Fig 2B**). Cells lacking both *ppx* and *ppk2* started accumulating polyP later than a *ppx* mutant alone but accumulated much more polyP at 48 hours and 72 hours than either the wild-type or the *ppx* mutant, while still having diminished polyP levels by 72 hours (**Fig 2B**), at which point almost all the cells had sporulated (see **Fig 4**). No polyP accumulated at any time point in a *spo0A* mutant (**Supplemental Fig 1**). Taken together, these findings support the idea that PPX is primarily responsible for degrading polyP, along with PPK2, and together they control levels of polyP accumulation within *B. cereus*, while PPK1 is primarily responsible for intracellular polyP synthesis in this species.

**Figure 2.**
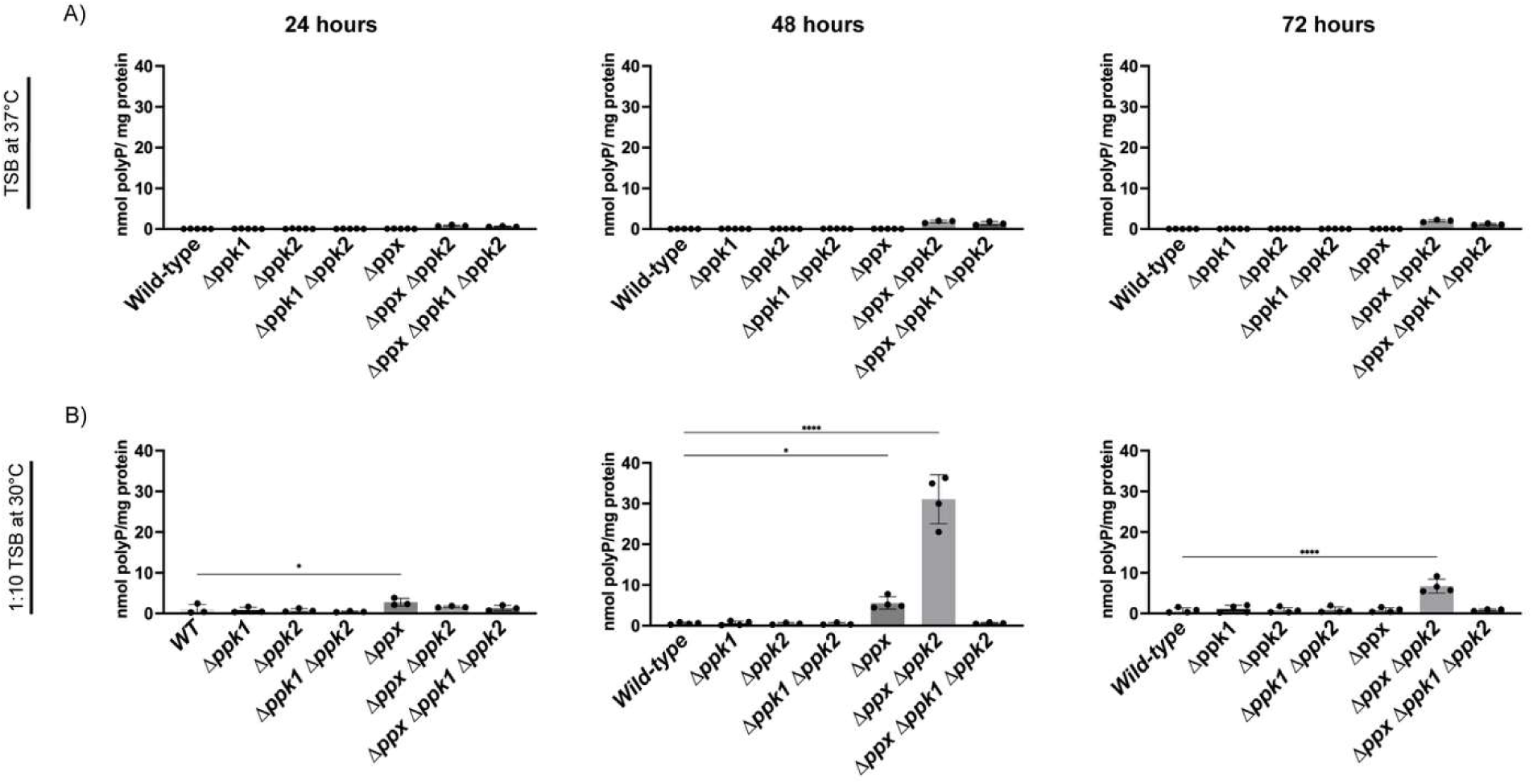
**A)** Accumulation of cell-associated polyP during growth of *B. cereus* in TSB at 37°C with shaking in beveled flasks over 72 hours **B)** Cell-associated polyP levels over 72 hours of cells grown in 1:10 TSB (sporulation media) at 30°C shaking in a beveled flask. Samples were taken over the course of 3 days. Statistics performed in Prism using a one-way ANOVA (* p=<0.05) (**** p=<0.0001).

### *B. cereus* produces very little polyP under traditional starvation conditions

PolyP accumulation is stimulated in *E. coli* and in *P. aeruginosa* by growing in rich medium (LB or TSB) and then shifting them to a minimal media such as M9 or MOPS minimal media ^41,70,79,85,86^. *E. coli* cells will accumulate nearly 200 nmol polyP/mg of protein by three hours after such a nutrient switch, then rapidly degrade that polyP ^39^. We wanted to test whether *B. cereus* cells would accumulate polyP in a similar manner to *E. coli* or *P. aeruginosa*. *B. cereus* cells were grown in TSB into exponential phase (OD_600_ = 0.2), before being rinsed and subcultured in MOPS minimal media. Cells experienced a short diauxic shift before resuming exponential growth (**Fig 3A**). We observed little to no polyP accumulation within the cell over the first 6 hours after switching to MOPS minimal media (**Fig 3B**), consistent with previous reports ^69^. We continued monitoring polyP levels within *B. cereus* over a period of several days and were observed a modest but reproducible spike of polyP at 24 hours of growth in MOPS minimal media (**Fig 3B**), more consistent with polyP synthesis as a characteristic of entry into stationary phase than of response to nutrient limitation. These findings suggest a deviation in *B. cereus* from known polyP accumulation bacterial models, highlighting the existence of species-specific mechanisms governing polyP production and storage.

**Figure 3.**
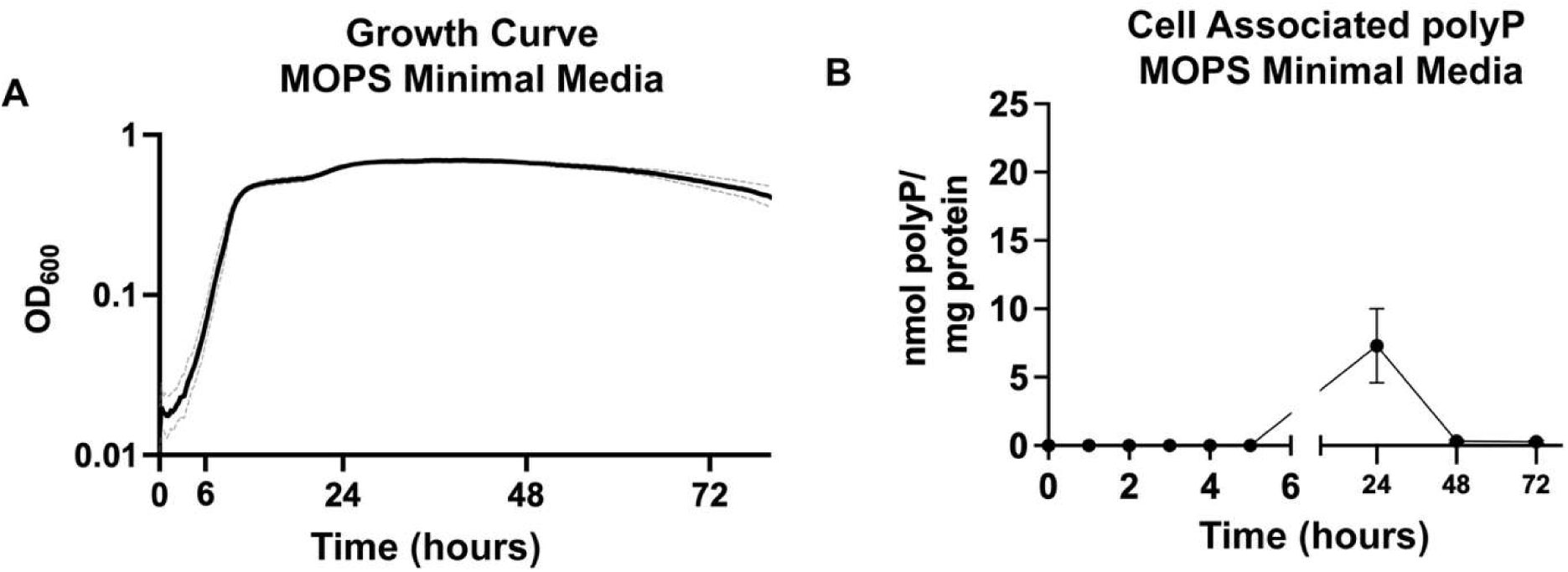
**A)** Growth of *Bacillus cereus* ATCC 14579 in MOPS minimal media (MOPS with Glutamic Acid 1.84 mg/ml, Leucine 0.8mg/ml, Valine 0.3 mg/ml, Threonine 0.168 mg/ml, Methionine 0.07 mg/ml, Histidine 0.05 mg/ml, 4 g liter^−1^ glucose, and 0.1 mM K_2_HPO_4_) at 37°C for 96 hours shaking. Culture density (OD_600_) was measured every 10 minutes and is indicated by solid black line, in triplicate. Error bars shown on graph as light gray dotted line. **B)** PolyP levels extracted from cultures at indicated times over the course of 72 hours.

### PolyP plays minimal roles in sporulation and motility

Previous work reported a modest reduction in sporulation efficiency in a *ppx* mutant ^69^. This observation is not entirely unexpected, given the established links between polyP metabolism, cellular metabolism, and cell cycle regulation in bacteria ^15,55^. Consistent with these findings, we observed a similar small reduction in sporulation in the *Δppx* mutant (**Fig 4A**), although it was variable enough that it was not statistically significant in our experiments. No other polyP mutants exhibited any defect in sporulation. As expected, a *spo0A* mutant made no spores (not shown)^84^. For unknown reasons, possibly relating to differences in strain or culture conditions, we were unable to replicate the reported motility defect of *B. cereus ppk1* mutants ^69^, although we did observe a substantial motility defect in the *spo0A* mutant (**Supplemental Fig 2**).

**Figure 4.**
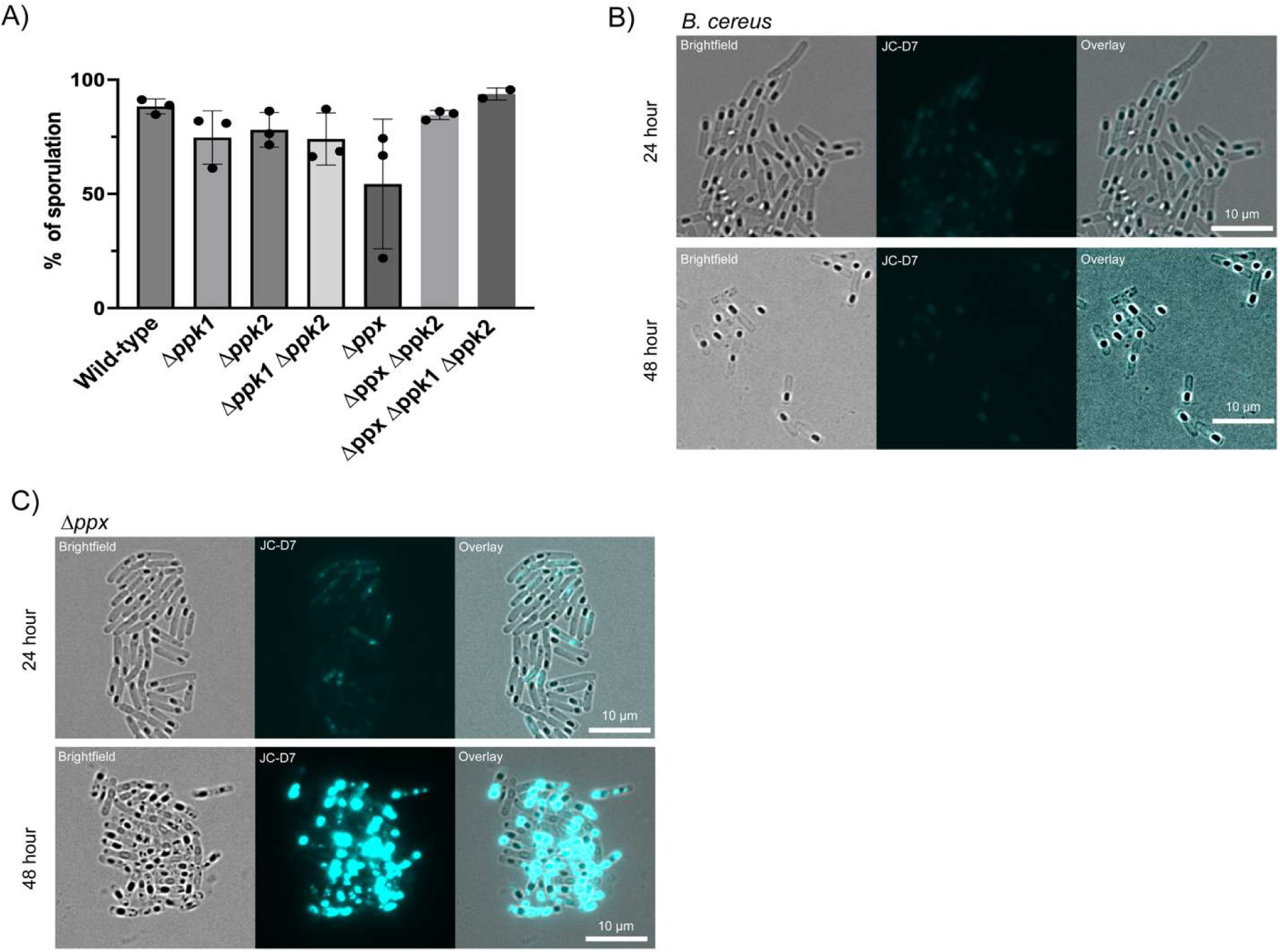
**A)** Percentage of cell populations with spores after 48 hours of growth in sporulation media at 30°C. By 72 hours all populations are ∼99% spores. **B)** Microscopy of wild-type *B. cereus* and **C)** *B. cereus Δppx* cells grown in sporulation media at 30°C in a beveled flask shaking. Cells were mixed with JC-D7 dye and spotted on agarose pads for imaging. JC-D7 was imaged using a custom JC-D7 filter cube (see Methods).

We next investigated whether intracellular polyP localization corresponded to spore formation by staining sporulating cells with the polyP-specific fluorophore JC-D7 ^76,77^ and visualizing polyP distribution. In wild-type cells, which contained little detectible intracellular polyP, only a weak diffuse signal was observed throughout the cell, with no apparent localization at either 24 or 48 hours of growth in sporulation medium (1:10 TSB) (**Fig 4B**). In contrast, the *Δppx* mutant, which accumulated an average of ∼5 nmol of intracellular polyP / mg total protein by 48 hours (**Fig 2B**), showed little detectible signal at 24 hours but exhibited intensely stained cytoplasmic polyP accumulation at 48 hours (**Fig 4C**). Staining with JC-D7 dye was sensitive enough that the signal from this small amount of polyP within the cells was enough to saturate our camera signal (**Fig 4C**). Overlay of the brightfield and JC-D7 fluorescence images revealed that these polyP accumulations were localized outside of the developing spore compartment, while still within the cell (**Fig 4C**). Taken together, these findings suggest that although polyP may influence the timing of sporulation, it is either not directly incorporated into or concentrated within the spore itself or the JC-D7 dye is unable to penetrate into spores. Regardless, these results show that micrographic staining with JC-D7 is easily able to detect levels of polyP in *B. cereus* near or below the limit of detection of our bulk culture quantification method.

### PolyP condensates within *B. cereus*

A recurring but unexplained feature of sporulating *B. cereus* is the presence of distinct intracellular granules that appear during spore development ^4,87^. Although these structures have been reported since the earliest studies of sporulation, their composition and function have remained unclear. We observed similar granules in stationary phase *B. cereus* cells (**Fig 5A**), and notably, all detectable granules exhibited JC-D7 fluorescence, indicating the presence of polyP (**Fig 5B-D**). We were puzzled to observe, however, that these JC-D7 bright granules were also present in both Δ*ppk1* and Δ*ppk1 Δppk2* mutants (**Fig 5C and 5D**).

**Figure 5.**
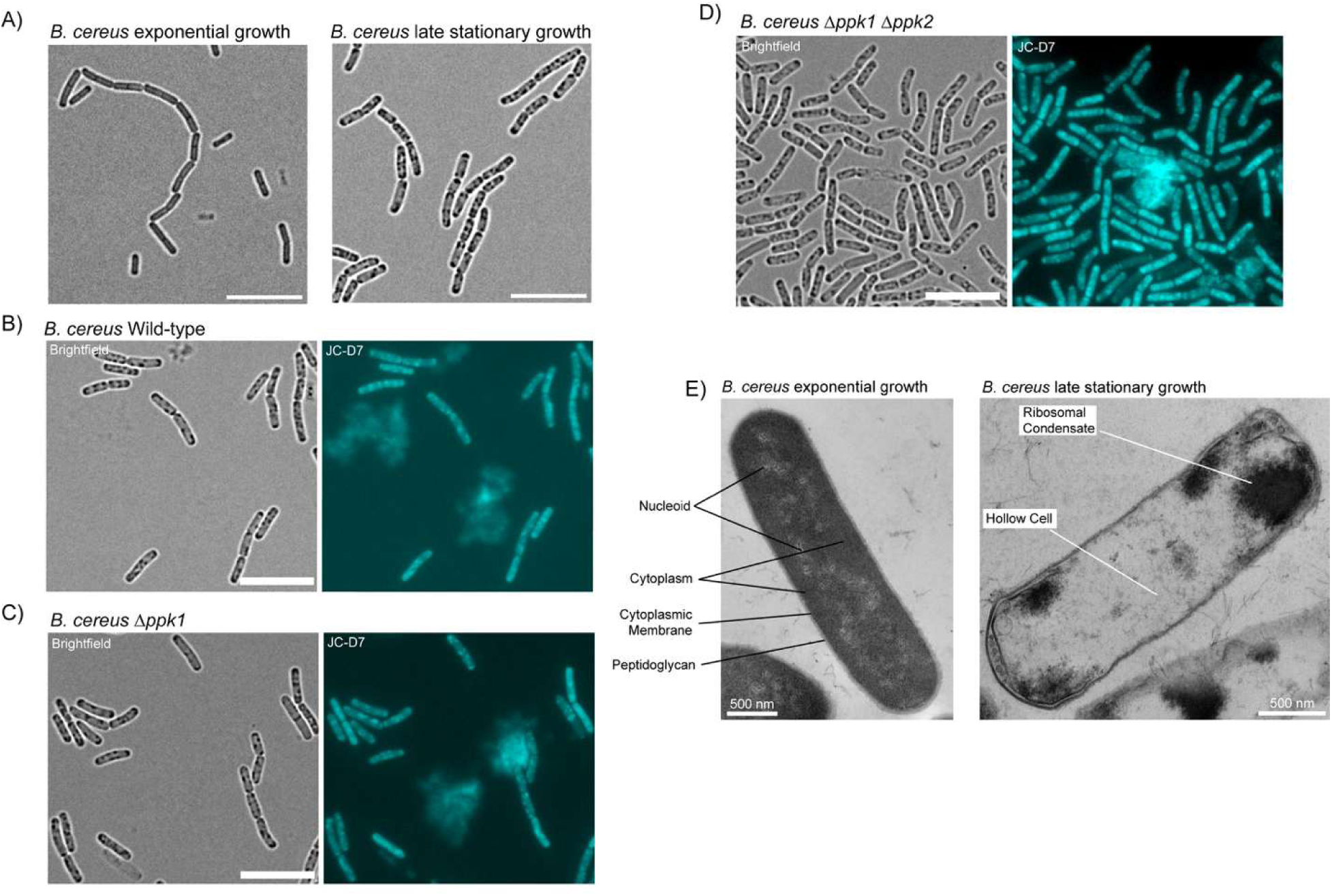
**A)** Brightfield microscopy of *B. cereus* wild-type cells at exponential phase of growth (3 hours) and at late stationary phase of growth (48 hours) in TSB media at 37°C. Unlabeled scale bars represent 10 µm. **B)** Brightfield and fluorescence microscopy with JC-D7 of *B. cereus* wild-type at late stationary phase of growth (48 hours) in TSB at 37°C**. C)** Brightfield and fluorescence microscopy with JC-D7 of *B. cereus Δppk1* at late stationary phase of growth (48 hours) in TSB at 37°C. **D)** Brightfield and fluorescence microscopy with JC-D7 of *B. cereus Δppk1 Δppk2* at late stationary phase of growth (48 hours) in TSB at 37°C. **E)** Transmission electron microscopy (TEM) of exponential phase growth (3 hours) and of stage stationary growth (48 hours). Scale bars are 500 nm for TEM images.

To determine whether these structures resembled the liquid-liquid phase separated polyP granules described in *P. aeruginosa* and other bacteria ^31,70^, we examined *B. cereus* cells by transmission electron microscopy (TEM) at 48 hours, corresponding to a time point at which intracellular JC-D7 positive granules are readily observed (**Fig 5A-D)**. Unlike the spherical, phase-separated polyP granules in *P. aeruginosa*, the structures observed in *B. cereus* appear as electron dense condensates within regions of the cytoplasm that otherwise appear depleted of ribosomes. Actively-growing cells without granules appeared as expected, with homogenously distributed ribosomes throughout their cytoplasm (**Fig 5E**). PolyP is known to interact with ribosomes ^88^, and the dimensions of the granular structures, approximately 200-500 nm in diameter, are consistent with the intracellular foci observed by light microscopy. Taken together, these observations suggest that the JC-D7 positive granules observed in *B. cereus* may represent ribosome condensates that form during nutrient limitation.

To investigate whether the presence of these granules was associated with altered cellular physiology, we performed time-lapse microscopy of wild-type cells from 48-hour cultures transferred to fresh TSB agarose pads and incubated at 37°C. Cells containing visible granules consistently failed to resume growth, whereas neighboring cells lacking granules readily initiated cell division (**Fig 6** and **Supplemental video 1**). Notably, granule containing cells remained growth-arrested throughout the entire 8-hour observation period despite exposure to nutrient rich conditions (**Fig 6**). To check and see if these cells were alive or dead we stained cells with propidium iodide, a common viability stain ^89^, and found that while the dye entered into some cells, indicating disrupted membrane permeability in those cells, not all the cells with granules inside allowed propidium iodide to pass (**Supplemental Fig 3**). This suggests that while some cells may have compromised membranes, many of these granule-containing cells are still alive but not growing.

**Figure 6.**
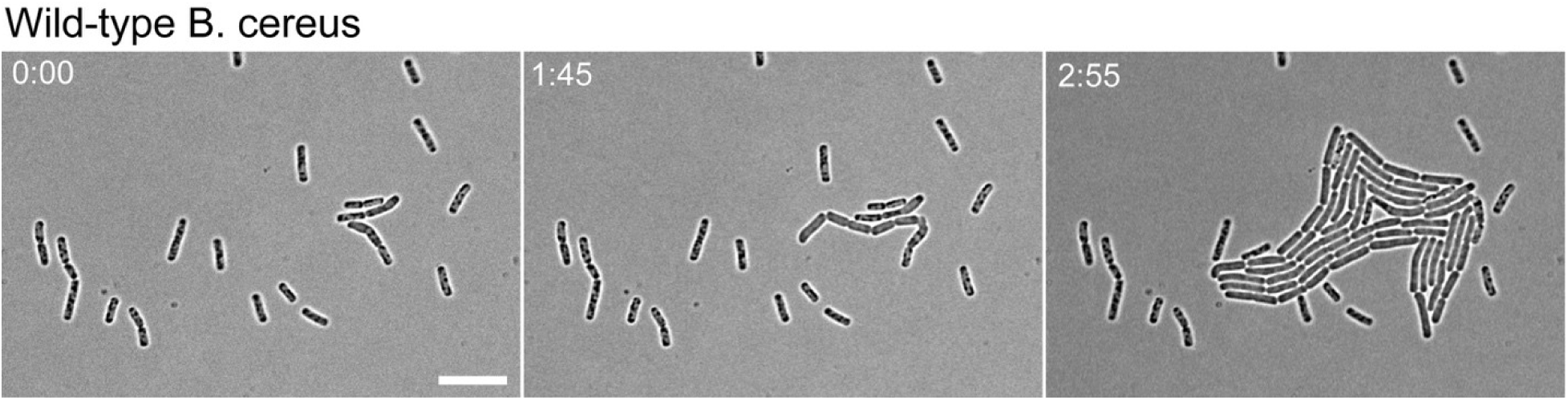
Time-lapse microscopy of wild-type *B. cereus* and *Δppk1* mutant strain. Strains were grown for 48 hours before being spotted on fresh TSB agarose pads and incubated at 37°C during time-lapse microscopy. Scale bars represent 10 µm, time is in hours and minutes.

### PolyP can be found outside the cell in *B. cereus*

Given the low levels of intracellular polyP that we detected, it was unclear whether *B. cereus* produced only limited amounts of polyP or whether polyP accumulated in a cellular compartment that is not typically examined. When imaging *B. cereus* cultures stained with JC-D7, in addition to discrete intracellular fluorescent foci, we frequently observed diffuse extracellular fluorescence surrounding subsets of cells (**Fig 5B and 5C**). These extracellular “clouds” were absent when regions of the agarose pad lacking cells were imaged, indicating that the signal originated from the cells or their surrounding medium rather than from background fluorescence.

Most studies of bacterial polyP focus exclusively on intracellular stores, leaving the presence and potential role of extracellular polyP largely unexplored. We therefore analyzed cell-free supernatants from growing cultures in TSB media at 37°C. PolyP release by the wild-type was minimal during early time points but became detectable by 24 hours and continued to increase over three days (**Fig 7A**). Under these conditions, extracellular polyP accumulated during late stationary phase, reaching up to ∼90 µM (**Fig 3**), while intracellular levels remained low (**Fig 2A**). Extracellular polyP accumulation was dependent on sufficient oxygenation (**Supplemental Fig 4**), with more highly aerated conditions promoting production, in contrast to *E. coli,* in which polyP accumulation is favored under anaerobic conditions ^90^. Most startlingly, the production of extracellular polyP was the same in all of the mutant strains, including the Δ*ppk1* Δ*ppk2* mutant, which lacks all known polyP synthesis enzymes (**Fig 7A**). This, in combination with the observation of JC-D7 bright granules in the Δ*ppk1* Δ*ppk2* mutant above (**Fig 5D**), led us to more seriously consider the possibility that in addition to PPK1 and PPK2, *B. cereus* encodes a third, previously unknown polyP synthesis pathway. Although JC-D7 fluorescence is thought to be specific to polyP ^76,77^, we used an established polyP-specific gel electrophoresis DAPI photobleaching technique ^82^ to independently confirm that the extracellular signal was due to the presence of *bona fide* polyP (**Fig 7B**). These findings indicate that extracellular polyP accumulation in *B. cereus* does not require the canonical polyP synthesis pathways. The observation that extracellular polyP production is independent of known polyP-synthesizing enzymes suggests the presence of previously uncharacterized mechanisms in the *B. cereus* group. Identification of the factors underlying this process will require further investigation.

**Figure 7.**
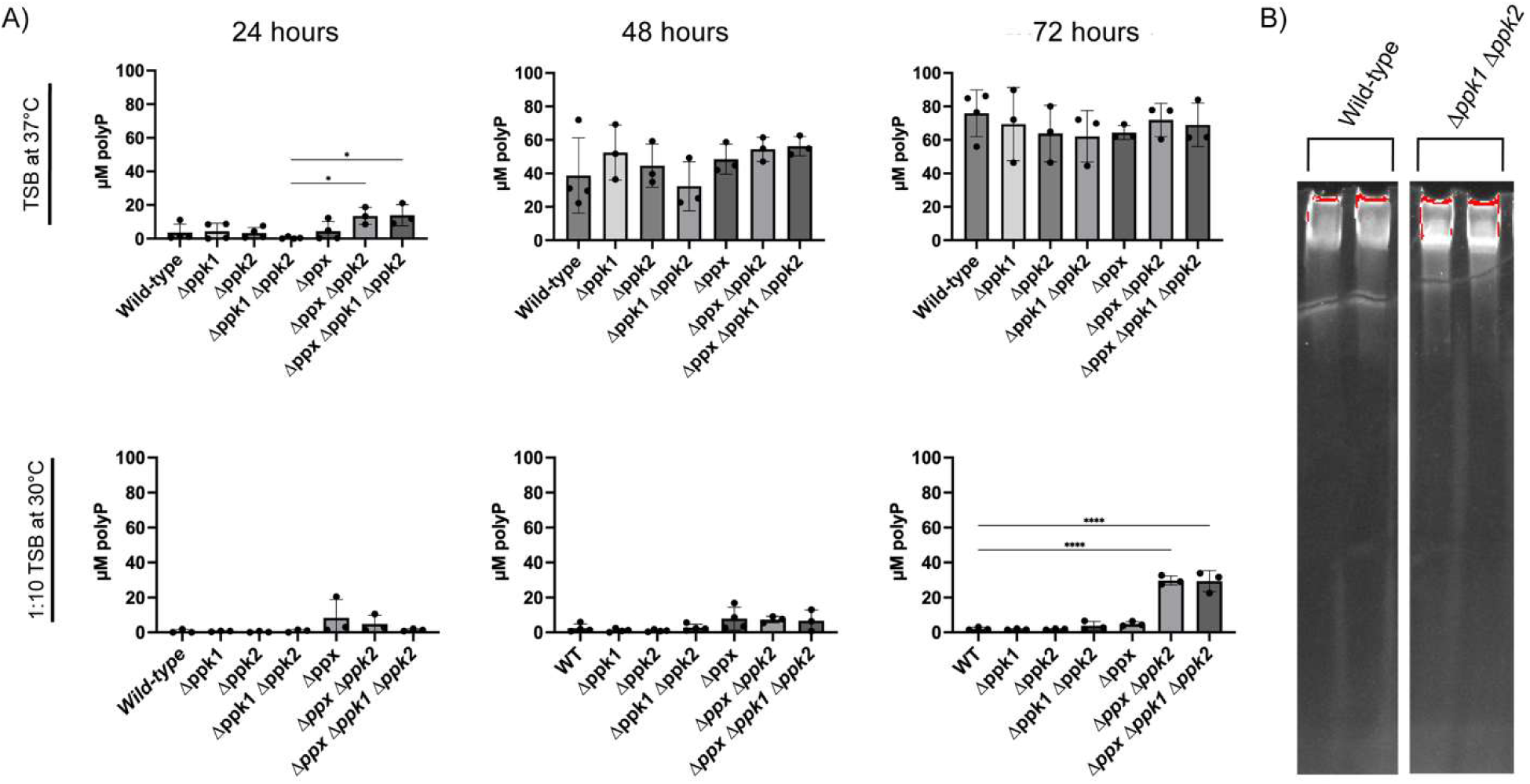
**A)** Accumulation of extracellular polyP in cell-free supernatants during growth of *B. cereus* in TSB at 37°C, or in1:10 TSB (sporulation media) at 30°C, with shaking in beveled flasks over 72 hours. Samples were taken over the course of 3 days. Statistics performed in Prism using a one-way ANOVA (* p=<0.05) (**** p=<0.0001). **B)** Aliquots of supernatants from strains grown for 72 hours at 37°C in TSB media shaking in a beveled flask. Supernatants were filtered through a 0.22 µm syringe filter, with polyP being extracted by phenol: chloroform extraction and separate on a 15% acrylamide TBE-urea gels and negative stained for polyP with DAPI.

We next examined if polyP was found extracellularly in cells during starvation as well and found that extracellular polyP levels remained low during the first 24 hours of growth in sporulation medium but increased modestly by 48 hours (**Fig 7B)**. After 72 hours, extracellular concentrations of polyP approached 30 µM in the *Δppx Δppk2* and *Δppx Δppk1 Δppk2* strains (**Fig 7B**). This indicated that extracellular polyP was PPK-independent under these conditions as well, but that, unlike in full-strength TSB, the polyP-degrading enzymes PPX and PPK2 did influence polyP release in 1/10^th^ TSB. No extracellular polyP was produced by the *spo0A* mutant under these conditions (**Supplemental Fig 5**). We also detected extracellular polyP accumulating in the supernatant of wild-type cells grown in MOPS minimal medium over the course of 72 hours (**Supplemental Fig 6)**.

### Extracellular polyP is resistant to ScPPX-mediated degradation

A common approach for detecting and quantifying polyP in prokaryotes relies on the use of purified ScPPX from *Saccharomyces cerevisiae* ^39,78,79^. To further characterize the extracellular polyP detected in culture supernatants, we treated samples with purified ScPPX. Surprisingly, no detectable hydrolysis was observed under conditions that readily degrade purified polyP controls (**Supplemental Fig 7**). ScPPX hydrolyzes polyP processively by removing phosphate residues from the terminal phosphoanhydride bonds of the polymer and therefore require access to the ends of the polyP chain ^65,78,91^. The resistance of the extracellular material to ScPPX-mediated hydrolysis suggests that the polymer may be inaccessible to the enzyme. Interestingly, polyP from the supernatant was still resistant to ScPPX degradation even after phenol/chloroform and ethanol precipitation (**Supplemental Fig 8)**. One possible explanation is that extracellular polyP is associated with proteins or other macromolecules that shield the terminal ends of the polymer and prevent enzymatic degradation, although phenol/chloroform extraction would be expected to remove most proteins. Alternatively, polyP may be incorporated into a larger extracellular structure that limits enzyme accessibility or the ends of the extracellular polyP chains might be covalently modified in a way that prevents ScPPX activity. Further studies will be required to determine the molecular basis of this resistance to ScPPX hydrolysis.

### Extracellular polyP is found in the related strains *B. thuringiensis* and *B. anthracis*

To investigate if this extracellular polyP was unique to *B. cereus* or if it was more widespread, we next examined the closely related species *Bacillus thuringiensis*, producer of insect-killing toxins, and a strain of *Bacillus anthracis* which is non-pathogenic to humans due to the lack of the pXO2 virulence plasmid ^72,92^. Extracellular polyP accumulation was observed in both *B. thuringiensis* and *B. anthracis* under similar conditions (**Fig 8**) suggesting that this phenotype is conserved across the *B. cereus* group and could have broader relevance during infection. Further studies need to be done in other pathogenic bacteria, both Gram-negatives and Gram-positives, to determine if extracellular polyP accumulation is widespread among bacteria and if it plays a role during infection by pathogenic bacteria.

**Figure 8.**
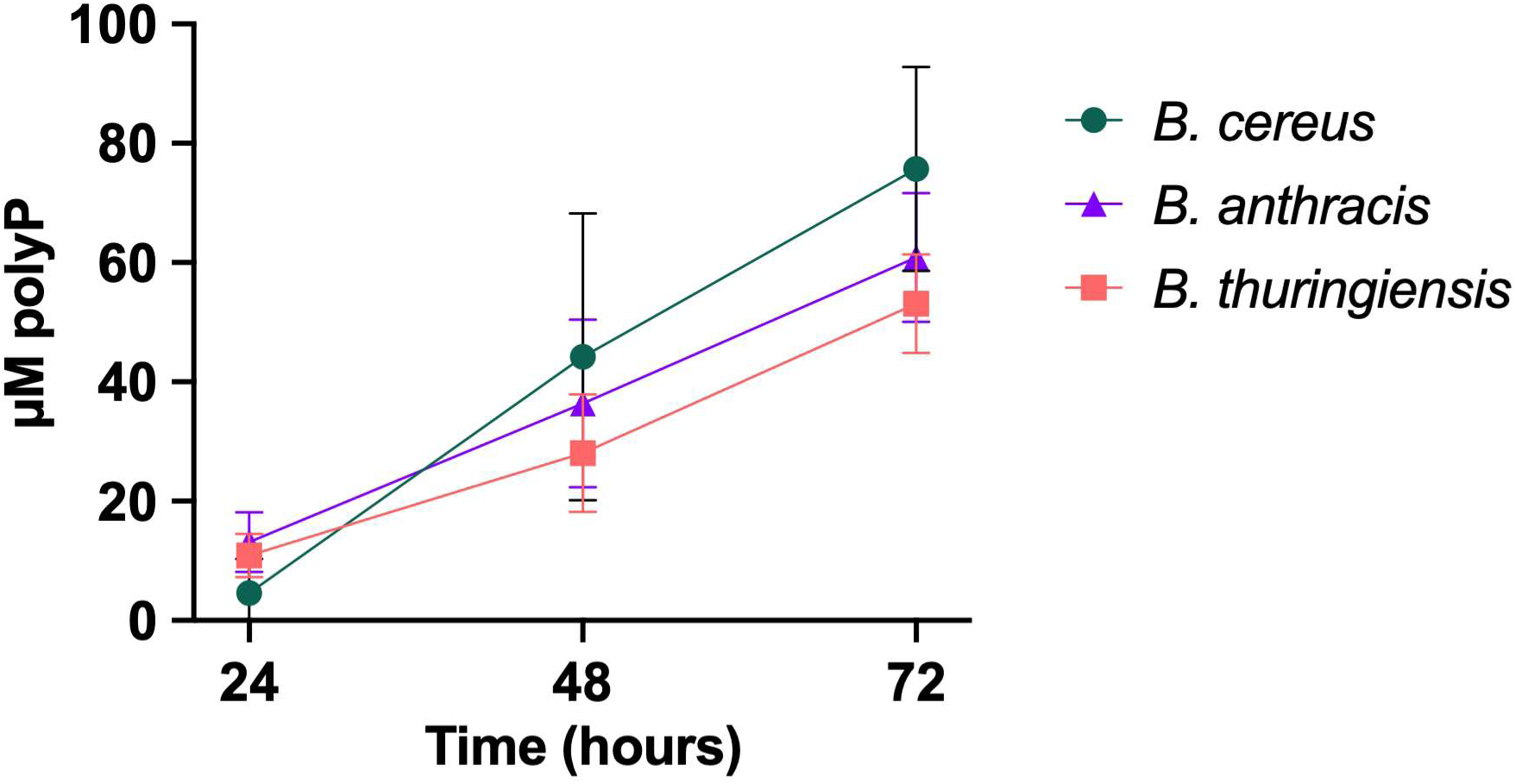
Accumulation of extracellular polyP in cell-free supernatants during growth of *B. cereus*, *B. thuringiensis* and *B. anthracis* in TSB at 37°C with shaking in beveled flasks over 72 hours.

## Discussion

Nearly all studies investigating bacterial polyP metabolism have been conducted in Gram-negative species ^14,30,31,33,38,43,44,50,55,70^. One notable exception is a single pioneering 2004 paper from Arthur Kornberg’s lab, which established *B. cereus* as a model organism for studying polyP biology in Gram-positive organisms ^69^. Despite this foundation, relatively little progress has been made in understanding the role of polyP in Gram-positive organisms over the past two decades. In the present study, we revisited *B. cereus* using contemporary approaches for polyP detection, microscopy, and genetic manipulation to investigate the functions of polyP throughout the bacterial lifecycle. Our findings reveal previously unrecognized roles for polyP during vegetative growth and sporulation, provide evidence for the existence of extracellular polyP associated with *B. cereus*, and most surprisingly, indicate the presence of a novel, non-PPK-driven mechanism for polyP synthesis. In fungi and some protozoa, polyP is synthesis is mediated by the vacuolar transporter chaperone complex component VTC4, encoding a transmembrane polyphosphate polymerase ^93,94^. No VTC4 homolog is present in *B. cereus*. The precise origin of *B. cereus* extracellular polyP remains unclear and identifying the protein(s) involved is currently a major focus of research in our lab.

PolyP has long been linked to bacterial survival, stress adaptation, and virulence during infection ^14,19,20,30,43,46,50,53,58,62,95,96^. More recent studies have demonstrated that bacterial polyP can accumulate at infection sites, inhibiting macrophage-mediated phagocytosis ^97^, modulate the immune response ^18^, and contribute to biofilm formation ^43,46,98^. However, the source of extracellular polyP present during infection has remained unclear. Our results here provide evidence that *B. cereus* cells can actively release or secrete polyP into the extracellular environment, suggesting a potential mechanism by which polyP accumulates at sites of infection and influences host-pathogen interactions. We further observed extracellular polyP in *B. thuringiensis* and *B. anthracis*, suggesting that this phenomenon is not unique to *B. cereus* and may be conserved among members of the *B. cereus* group and, potentially, other Gram-positive organisms. While the broader distribution of extracellular polyP among bacteria remains unknown, these findings raise the possibility that extracellular polyP may be more widespread than currently appreciated and could contribute to host-pathogen interactions in other organisms. These observations also raise questions regarding the mechanism by which polyP reaches the extracellular environment. For example, extracellular polyP could be released following cell lysis, actively exported from the cell, or secreted in association with proteins or other macromolecular assemblies. We are working to identify these mechanisms in *Bacillus* species now, although we expect that rigorous determination of polyP export mechanisms will not be possible until we identify the source of non-PPK polyP synthesis in these bacteria.

We observed the accumulation of intracellular granules prior to sporulation in starving cells (**Fig 5**). These granules have been reported previously ^4,87^, but their composition and biological significance remain unclear. The fact that these granules stain with JC-D7, suggesting they contain polyP even in the absence of PPK1 and PPK2, supports the idea that in *Bacillus* species there is a non-canonical polyP producing enzyme. One potential interpretation of these observations is that polyP participates in the formation of macromolecular assemblies containing ribosomal components, potentially contributing to a reduction in translational activity during entry into sporulation. Although polyP has long been known to associate with and modulate ribosomes ^88^, the functional significance of this interaction remains incompletely understood. Our findings are in agreement with the possibility that polyP may contribute to the regulation of translational activity under specific physiological conditions ^99–101^. Protein synthesis is an energetically demanding process that is central to bacterial growth and survival ^102–106^. To adapt to changing environmental conditions, bacterial cells tightly regulate protein synthesis, trafficking, folding and degradation ^103,107,108^. One strategy used by bacteria involves ribosome hibernation factors which promote the conversion of active 70S ribosomes in translationally inactive 100S ribosomes during stress ^108^. Additional mechanisms capable of reducing protein synthesis include ribosomal pausing or stalling on specific codons ^109^ or through mRNA secondary structures ^110^, so it is possible that these large ribosome complexes represent a new mechanism for ribosome inactivation. However, the composition of the JC-D7 bright dense granules in *B. cereus* and the extent to which they might directly influence ribosome function remain unknown.

A speculative extension of this model is that polyP accumulation may serve dual roles during periods of reduced growth in *B. cereus*. In addition to any potential regulatory effects on protein synthesis, polyP could function as a reservoir of stored phosphate and metabolic potential that can later be mobilized when favorable conditions return. Under such circumstances, polyP degradation could both relieve any polyP-associated inhibitory effects and provide phosphate for rapid metabolic reactivation. While this hypothesis is consistent with the established roles of polyP in cellular energetics, our current data do not directly address these mechanisms. Future studies aimed at defining the molecular composition of polyP containing granules, determining whether ribosomes are associated with these structures, and measuring translational activity in the presence and absence of polyP will be necessary to evaluate these possibilities and establish any mechanistic connection between polyP accumulation and translational regulation.

---

**Supplemental Figure 1.**
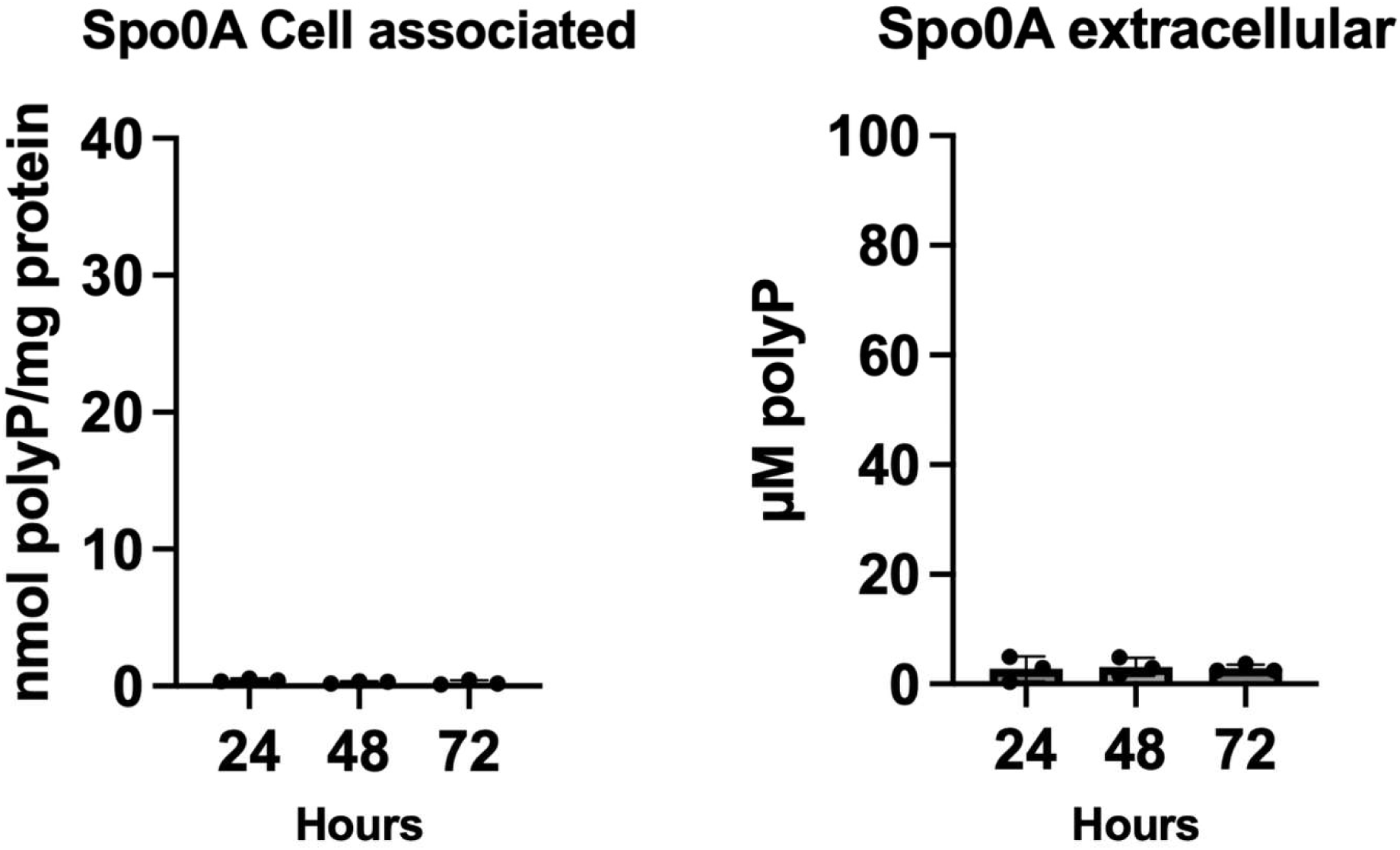
Analysis of both cell associated polyP levels over 72 hours of *B. cereus* Δ*spo0A* cells grown in 1:10^th^ TSB (sporulation media) at 30°C shaking in a beveled flask. Samples were taken over the course of 3 days. Statistics performed in Prism using a one-way ANOVA.

**Supplemental Figure 2.**
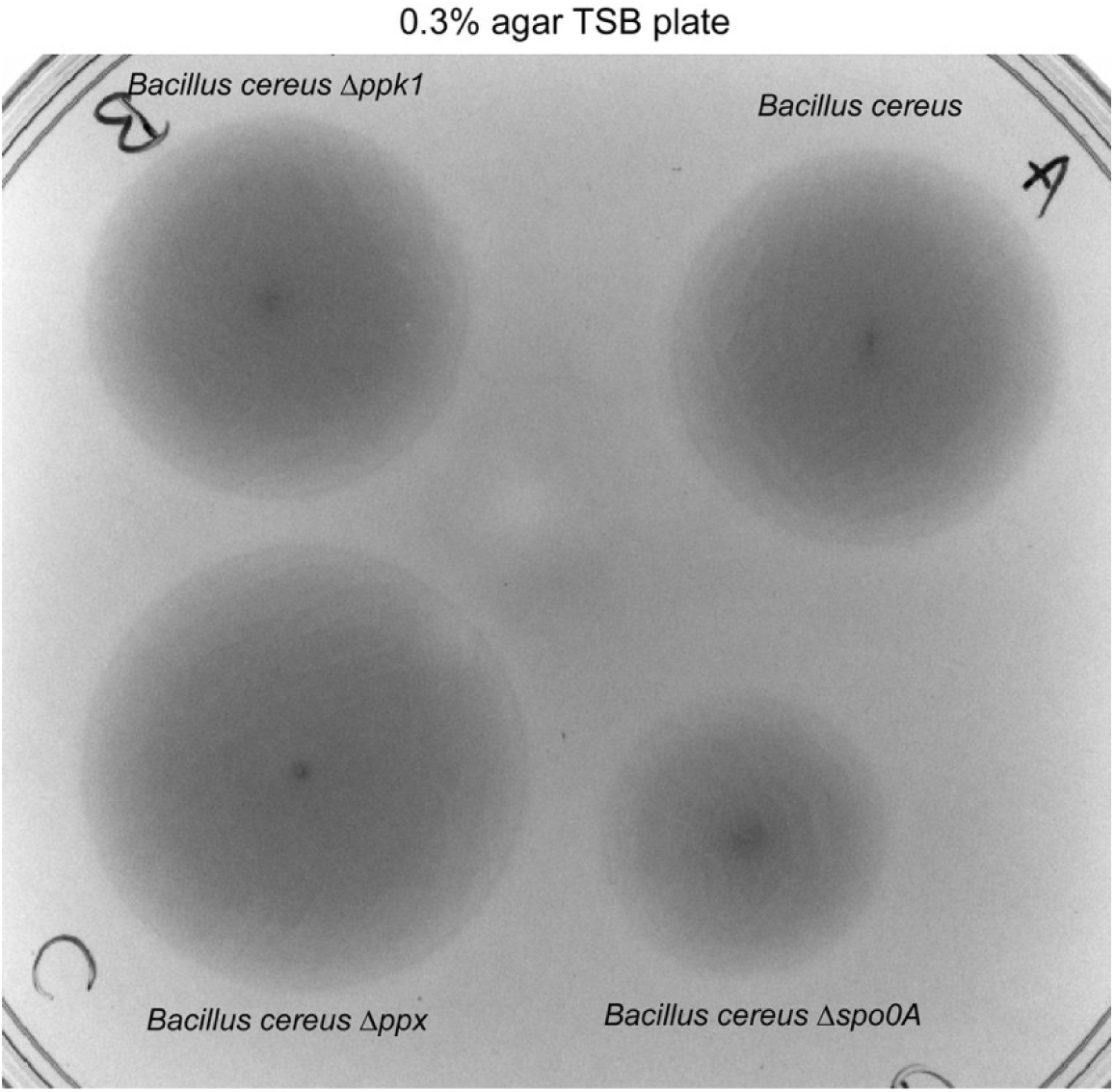
Cells were spotted on a 0.3% agar TSB plate, and allowed to grow at 37°C for 24 hours.

**Supplemental Figure 3.**
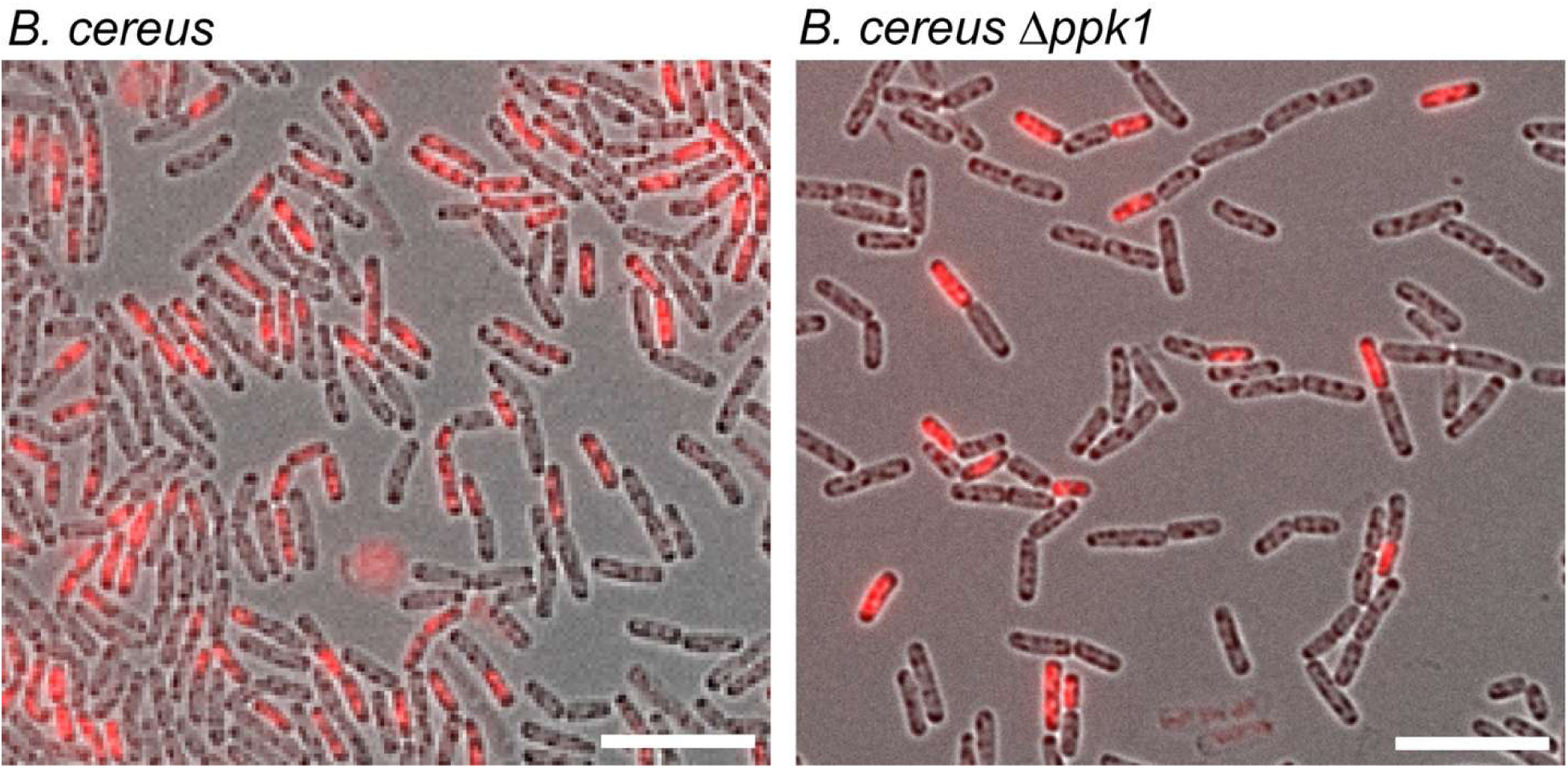
Microscopy of Wild-type *B. cereus ATCC 14579* (CH0082) and *Δppk1* mutant (CH0193) growth in TSB at 37°C for 48 hours spotted on agarose pads. Propidium iodide was added prior to imaging and images were captured in Brightfield and with a TritC filter cube.

**Supplemental Figure 4.**
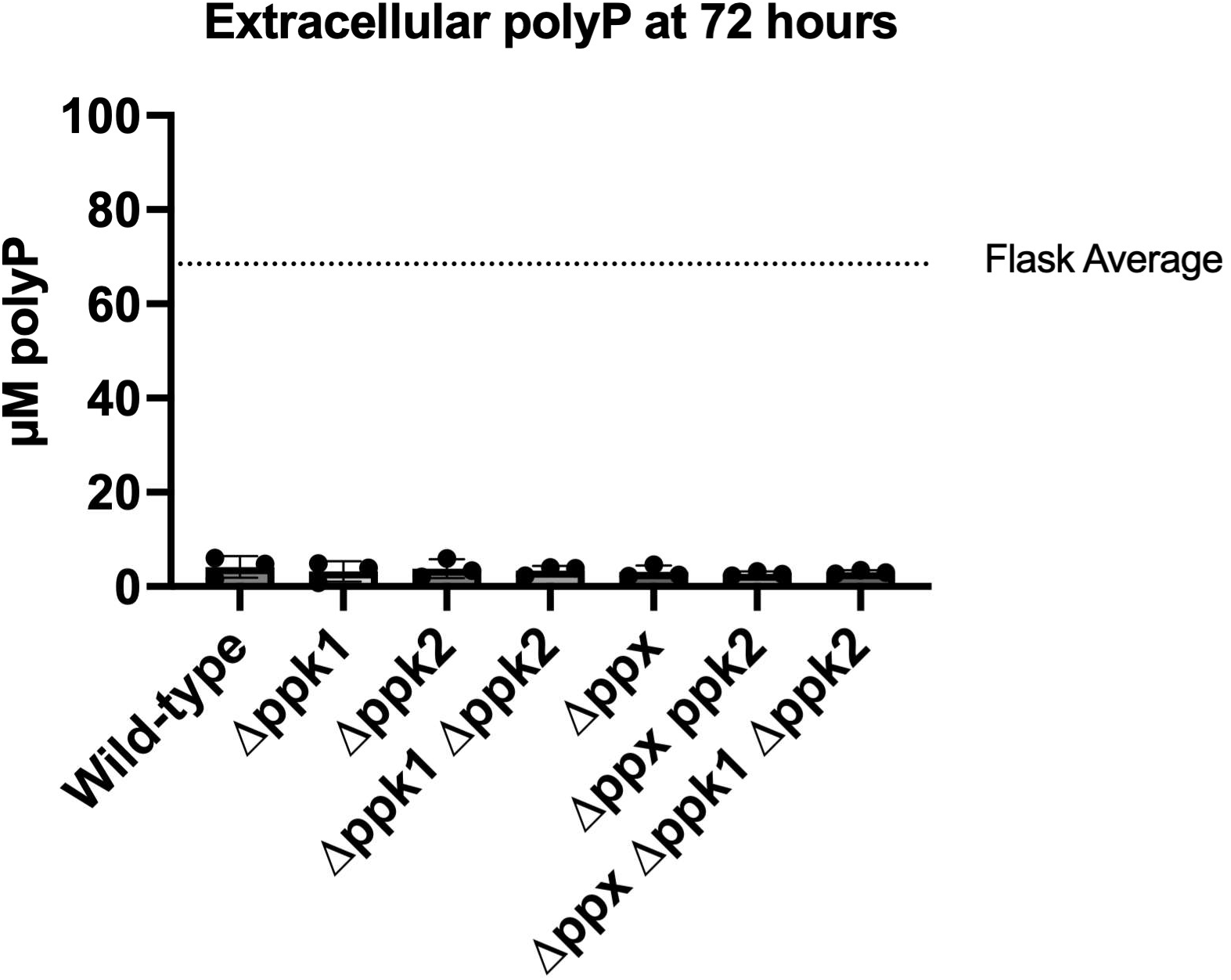
Extracellular amounts of polyP were quantified from cells grown in 5 ml TSB at 37°C shaking at 180-rpm in glass test tubes. Dotted line is average of all extracellular polyP values grown in TSB in beveled flasks from Figure 7A at 72 hours.

**Supplemental Figure 5.**
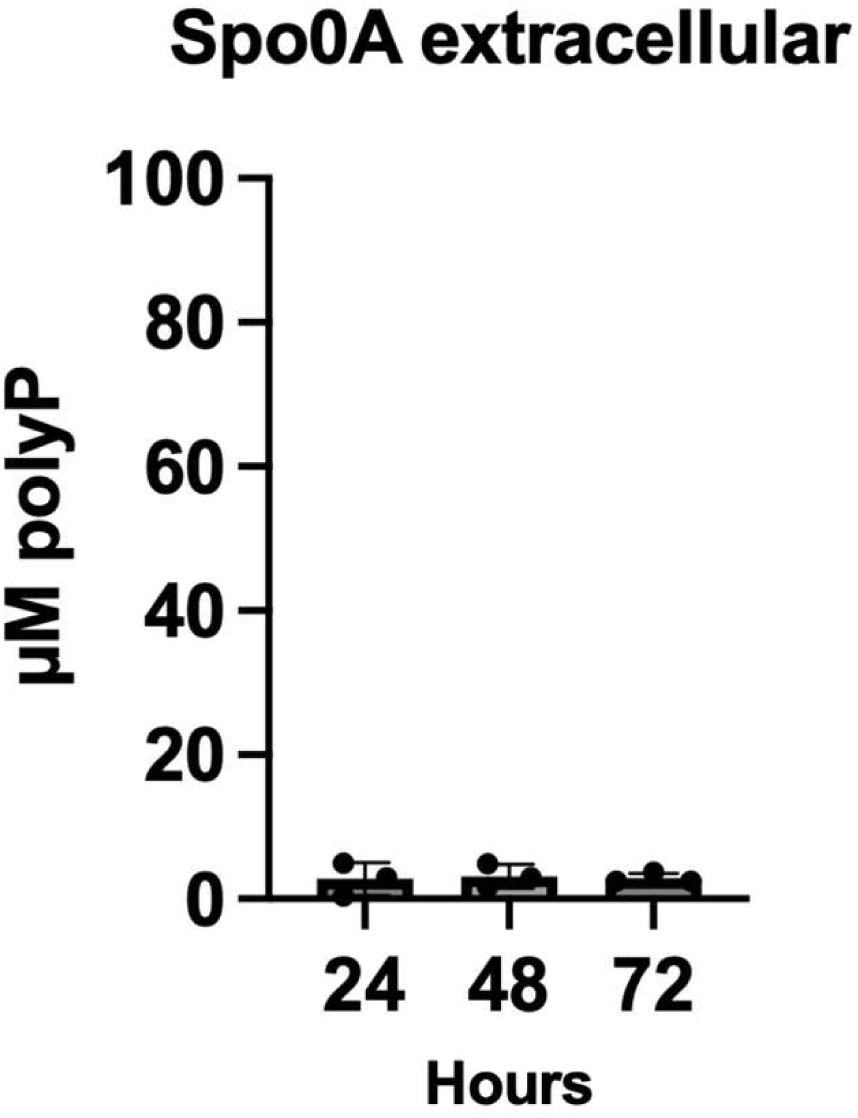
Analysis of extracellular polyP levels over 72 hours of *B. cereus* Δ*spo0A* cells grown in 1:10^th^ TSB (sporulation media) at 30°C shaking in a beveled flask. Samples were taken over the course of 3 days. Statistics performed in Prism using a one-way ANOVA.

**Supplemental Figure 6.**
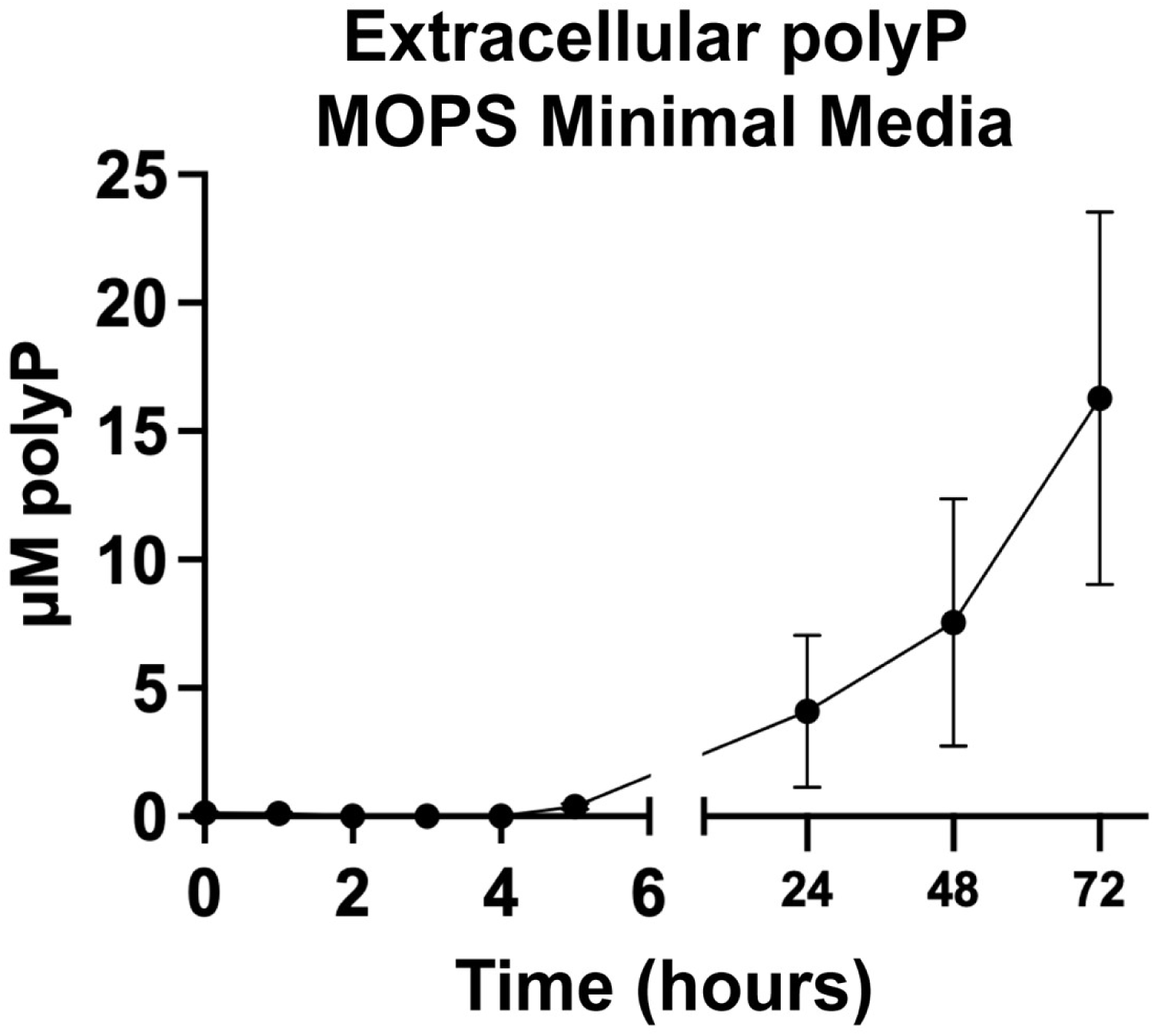
polyP levels found in the supernatant of growing cultures. Cells were grown in MOPS minimal media MOPS with Glutamic Acid 1.84 mg/ml, Leucine 0.8mg/ml, Valine 0.3 mg/ml, Threonine 0.168 mg/ml, Methionine 0.07 mg/ml, Histidine 0.05 mg/ml, 4 g liter^−1^ glucose, and 0.1 mM K_2_HPO_4_) at 37°C shaking in beveled flasks and matched samples for B and C were taken at indicated times in triplicate.

**Supplemental Figure 7.**
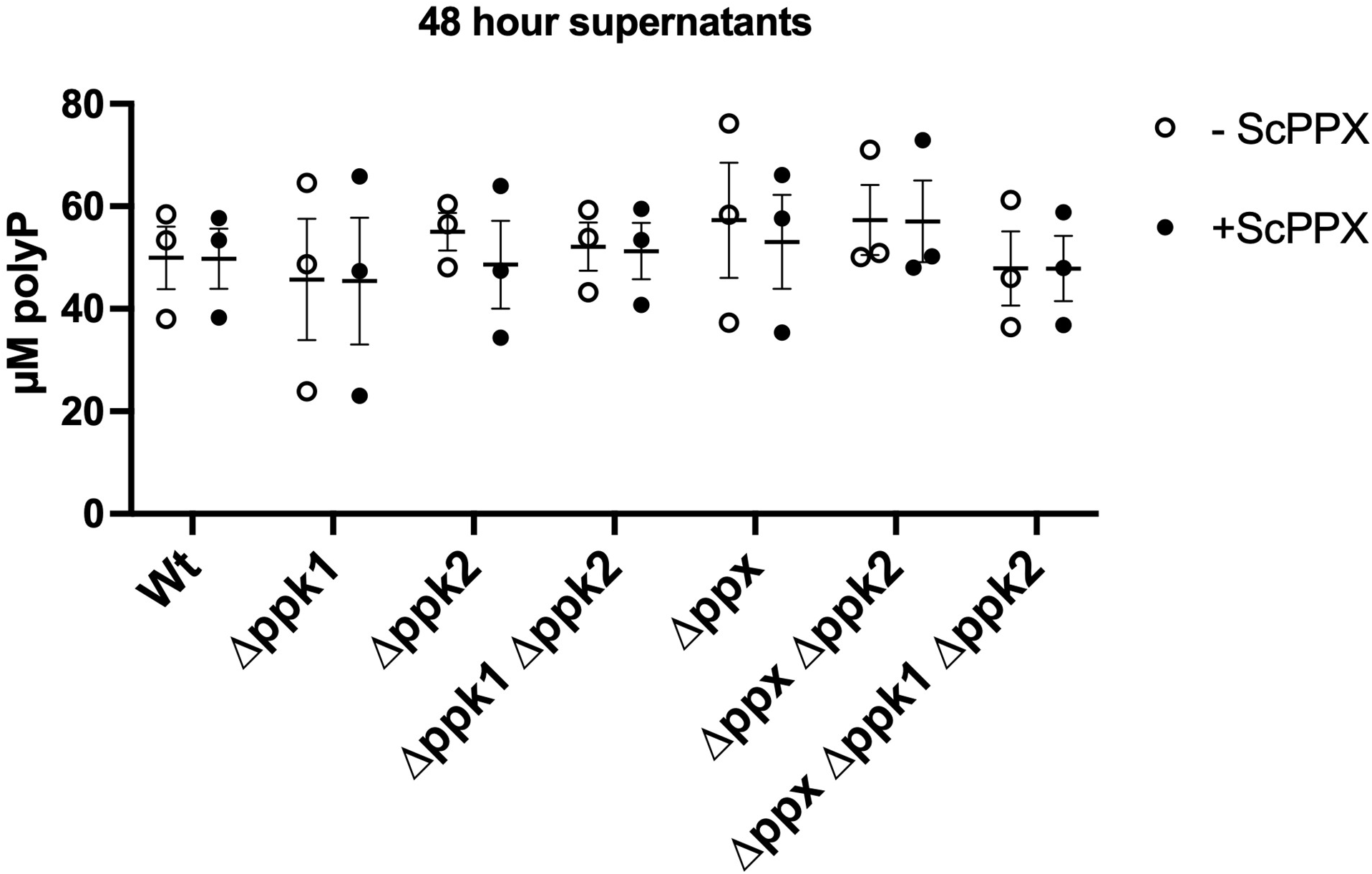
ScPPX degradation of 48-hour supernatants of cells grown in TSB at 37°C for 48 hours. Supernatants were treated with ScPPX and incubated for 20 minutes at 37°C.

**Supplemental Figure 8.**
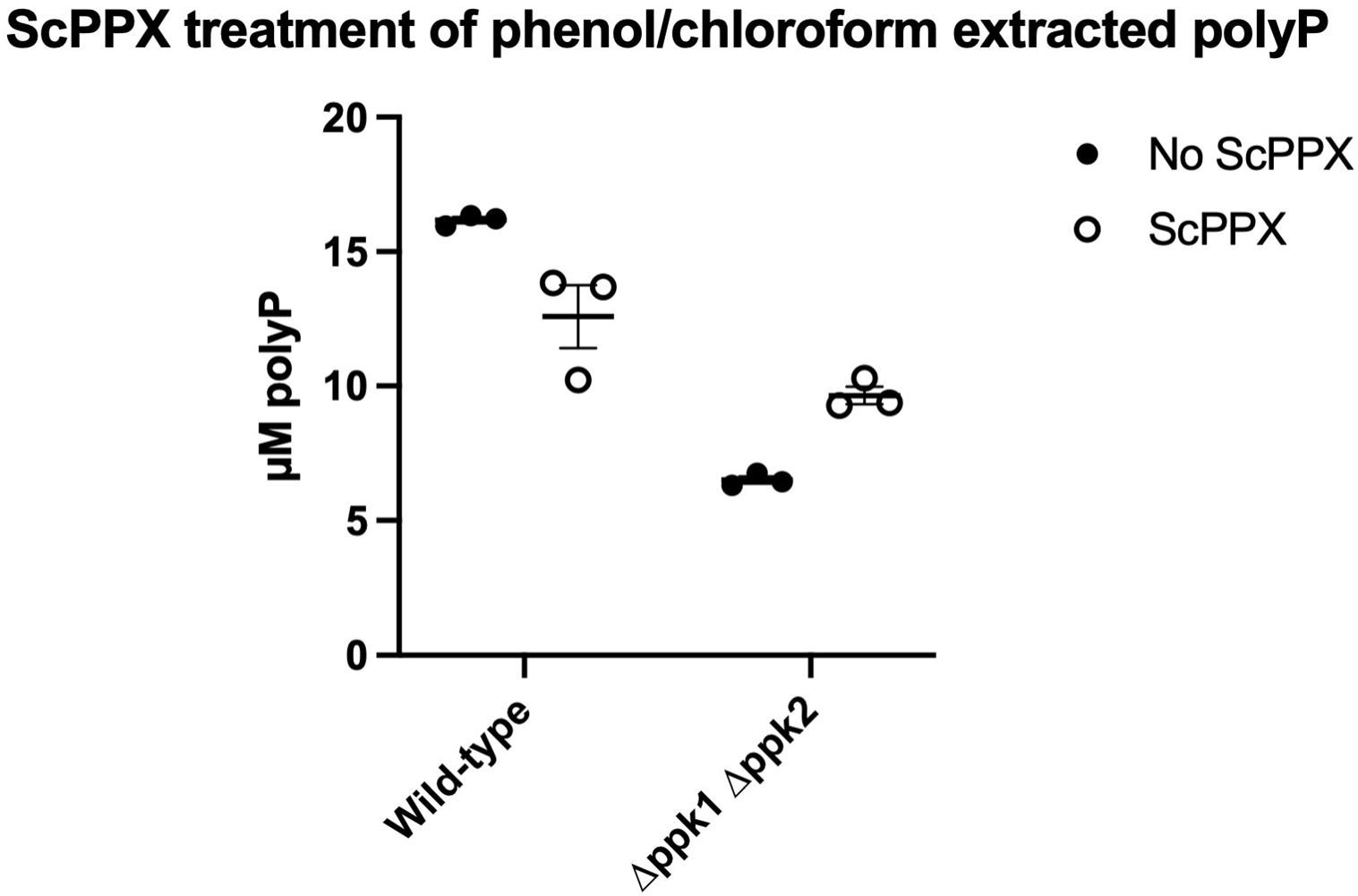
ScPPX degradation of phenol/chloroform extracted and ethanol precipitated polyP from supernatants of cells of the indicated genotypes grown in TSB at 37°C for 72 hours. Resuspended polyP solutions were treated with ScPPX and incubated for 20 minutes at 37°C. These pools are matched with the samples from Fig 7B, with 5 µl of extracted sample being mixed with 145 µl of 25 mM HEPES-KOH pH 8.0 prior to adding ScPPX buffer and enzyme for degradation.

Supplemental Video 1. Timelapse microscopy of a Wild-type *B. cereus* (CH0082) cells grown for 48 hours in TSB at 37°C spotted on a fresh TSB agarose pad, incubated at 37°C and imaged every 5 minutes.

## Acknowledgments

This work was funded by NIGMS R35 MIRA grant GM124590 (to MJG) and NIGMS F32 postdoctoral fellowship GM159427 (to CWH). Thank you to the Bacillus Genetic Stock center for kindly providing *Bacillus cereus* ATCC 14579 for this study.

